# Changes of platelet morphology, ultrastructure and function in patients with acute ischemic stroke based on super-resolution microscopy

**DOI:** 10.1101/2022.11.29.518340

**Authors:** Bingxin Yang, Xifeng Wang, Xiaoyu Hu, Yao Xiao, Xueyu Xu, Xiaomei Yu, Min Wang, Honglian Luo, Jun Li, Yan Ma, Wei Shen

**Affiliations:** Department of Neurology, Wuhan Puai Hospital, Tongji Medical College, Huazhong University of Science and Technology, Wuhan, Hubei 430030, China; Wuhan Blood Center-Huazhong University of Science and Technology United Hematology Optical Imaging Center, Hubei Institute of Blood Transfusion, Wuhan Blood Center, Wuhan, Hubei 430030, China

## Abstract

Platelets are important in acute ischemic stroke (AIS) thrombosis. The observation and evaluation of platelet ultrastructure and efficacy of antiplatelet drug in AIS patients is difficult due to microscopic limitations and sensitivity of platelet. The new super-resolution microscope (SIM) can accurately, quickly analyze the platelet ultrastructure and antiplatelet drug in AIS patients. We applied SIM to observe the morphology and ultrastructure of platelets with AIS patients in different state. SIM images were analyzed to specify the dense granules and α granules change of platelets in AIS patients. Testing platelet factor 4 (PF4) to reflect platelet releasing function. We observed that platelet activation in AIS patients was greater after stimulation, with α granule showing a pattern of parenchymatous masses. SIM images analyzing showed diameter of platelets, average size of granules, area% of granules per field and mean area of granules per platelet in AIS patient were lower than healthy people. Platelet releasing function was suppressed at rest stage and more efficacy release after stimulation. 2MeSamp inhibited parenchymatous masses of α granules and reduced PF4 release of platelets after stimulation. According to the results, the structure and function of platelets in AIS patients are indeed altered. Additionally, SIM could be used as a new method to indicates the onset of AIS and assess antiplatelet drugs.

## Introduction

Acute ischemic stroke (AIS) is the second leading cause of death worldwide, the most fatal sudden disease after heart disease, and the leading cause of disability [1, 2]. By 2030, stroke-related disability is projected to be the fourth most important cause of disability-adjusted life years in developed countries [3]. AIS refers to the sudden termination of blood supply to the brain tissue when the cerebral blood flow is below the ischemic threshold (10-25% of baseline)[4], causing irreversible nerve cell damage in the corresponding ischemic area and eventually lead to nerve function impairment. Intracranial atherosclerosis is the most common cause of AIS, and platelets play an inevitable role in subsequent vascular blockage[5].

Platelets are small anuclear cells, about 2 to 4μm in diameter[6]. Under physiological conditions, it plays the role of hemostasis, inflammation and immune response. On the contrary, it is overactivated to form thrombosis in some pathological situations such as ischemic stroke or acute myocardial infarction[7, 8]. In AIS patients, platelet surface is exposed to circulating thrombin and von willebrand factor (vwf) which elevated after atherosclerotic plaque ruptured and subsequently activated[9]. During platelet activation, platelets ultrastructure named dense granules and alpha granules can release a large number of granular contents. The dense granule release adenosine diphosphate (ADP), serotonin (5-HT) and calcium ions to recruit more platelets. And α granule expand the aggregation effect by releasing β-thromboglobulin (β-TG), platelet factor 4 (PF4) and a large number of coagulation factors[10]. With the participation of these granules, platelet throughout adhesion as well as aggregation, and eventually forms thrombosis to block the cerebral blood vessel, leading to the occurrence of AIS [2]. Some studies have pointed out that platelets are more likely to be activated with the increase of age, which may contribute to the higher incidence of AIS in elder people [11]. As highly reactive blood cells, platelets have undergone significant changes in their ultrastructure in some diseases, such as changes in platelet microtubules and mitochondria in patients with ovarian cancer [12]. Another study noted a significant reduction in the dense granules of platelets in patients with Hermansky-Pudlak syndrome [13]. In view of the tightly connect between AIS and platelets, studying the changes of platelet morphology, ultrastructure and granule function in AIS patients will be vital for the diagnosis and treatment of this disease.

Antiplatelet drugs are widely used for secondary prevention of AIS. According to the 2018 American Heart Association/American Stroke Association and 2019 BMJ Clinical Practice guidelines, early combined aspirin and clopidogrel should be used in patients with mild stroke and transient ischemic attacks (TIA), but long-term use is not recommended[14, 15]. As a thromboxane A2 (TXA2) inhibitor, aspirin irreversibly inhibiting platelet cyclooxygenase-1 (COX-1) to prevent the formation of TXA2[9]. Clopidogrel acts as an inhibitor of ADP receptor which activates platelets by binding to two protein receptors on platelets (P2Y1 and P2Y12) [16]. However, the majority (82 to 87%) of patients taking aspirin alone are not protected from further vascular events [17]. It also has been suggested that clopidogrel may have a benefit over aspirin in patients with ischemic cerebrovascular disease (CVD) [18]. Which drug is better for AIS patients is still controversial and it may need more detail study of platelets to support. Therefore, our study observed the alteration of the two drugs on the ultrastructure of activated platelet dense granule, α granule morphology and granular release function to compare the difference effect on platelet subcellular structure between the two drugs.

In most studies, platelet function was measured by light transmission aggregation assay (LTA), flow cytometry, thrombus elastography and other methods[19]. SIM as a new kind of optical imaging technology, with high resolution, rapidity and unbiased data, can provide convenient and reliable analysis of platelet ultrastructure. Existing research have applied SIM to investigate platelet ultrastructure in illness state, and provided new ideas for disease diagnosis[12, 13, 20]. Since the releasing of ultrastructural dense granules and α granules of platelets are critical in platelet activation of AIS patients, four different states of the two granules in AIS patients were observed and analyzed by SIM, including (1) rest; (2) activate; (3) activation after 2MeSAMP treatment; (4) activation after Aspirin treatment. 2MESAMP and Aspirin are two in vitro antiplatelet drugs that inhibit cyclooxygenase-1 (COX-1) and P2Y12 receptor, respectively. Ultrastructure observation and analysis of platelet by SIM may offer more details of platelets in AIS patients. Through the changes of platelet ultrastructure in the elderly can be monitored to predict the occurrence of this disease. Meanwhile, the reactivity of platelets to different antiplatelet drugs could be evaluated to supply reference for clinical individualized medication.

## Materials and methods

### Materials and reagents

Thrombin, Aspirin and 2-methyl-thioadenosine-monophosphate (2MeSAMP) were purchased from Sigma (America). We made a purchase of 8%Paraformaldehyde Aqueous Solutuon (PFA) from Electron Microscopy Sciences (America).Phosphate buffered saline (PBS)was purchased from Tianjin Haoyang Biological Manufacture Co.,Ltd (China).We bought Anti-CD63 (ab8219),Alexa Fluor dyes 488 (ab150113) and Alexa Fluor dyes 647 (ab150079) from abcam (England). Anti human Von Willebrand Factor (vwf) was purchased from Dako(Denmark).15ml and 50ml centrifuge tube were purchased from Nest Biotechnology Co.,Ltd (China).Glass bottom cell culture dish was purchased from Guangzhou saiguo biotech Co.,Ltd (China). Human platelet factor 4(PF4) ELISA Kit were purchased from Cusabio (China).

### Patients’ selection

Patients with a diagnosis of AIS according to Chinese guidelines for diagnosis and treatment of acute ischemic stroke 2018 were recruited from Wuhan Puai Hospital. Other inclusion criteria were: (1) over 18 years old without pregnancy or suckling; (2) the incidence time of ischemic stroke was 24 hours at the most; (3) all patients received computed tomography (CT)or magnetic resonance imaging (MRI) of brain and responsible focus have been observed; (4) focal neurological deficits as well as global neurological deficits attributed to cerebral ischemia. Exclusion criteria was made of: (1) cerebral hemorrhage or other pathology situations including cerebral vascular malformation, brain tumor and encephalopyosis were found in brain CT or MRI; (2) the patients who have taken any type of antiplatelet drugs or medicines influence the function of platelets in 14 days; (3) history of atrial fibrillation, severe hepatic insufficiency, end-stage renal disease, autoimmune diseases, malignant tumor, blood system disease; (4) gastrointestinal bleeding or major surgery during 3 months; (5) administration of intravenous anticoagulants or thrombolytics within 1 week; (6) having a fever or infectious disease in 2 weeks.

### Control group choosing

Age may influence the function and structure of platelets[21], consequently we choose control groups divided into young healthy (YH,18~35 years old) and elderly healthy (EH, over 60 years old) the same time in order to avoid age confounding factor. Control group came from neurology ward of Wuhan Puai Hospital from December 17, 2021 to June 21, 2022. Both of the controls need to meet these requirements that mentioned above except suffering an acute ischemic stroke.

### Platelet preparation

Samples were collected from December 17, 2021 to June 21, 2022. All the volunteers signed informed consent forms. We have acquired ethic approval through Wuhan Forth Hospital and the approval number is Lun Shen Zi(KY2022-086-01). Human whole blood was collected into a 1:9 solution of sodium citrate from neurology ward of Wuhan Puai Hospital. The collected blood was centrifuged at 200×g for 12 min in 6 hours to obtain platelet-rich plasma (PRP). After incubating in 37°C for 2 hours, the PRP was treated with either 2MeSAMP, a P2Y12 receptor antagonist of platelet, or aspirin which can inhibit COX-1. Concentration of 2MeSAMP is 10uM, and 53uM of aspirin. Then, the platelet added in antagonist was co-incubated in 37°C for 10 min. Adding 0.5U thrombin to activate the platelet for 5 min or 1 hour. The supernatant of activated platelet was collected for ELISA test. 8%PFA in PHEM at a final concentration of 4%PFA was used to fix the remaining platelet for 15 min. Washed with PBS for 3 times, permeabilization with 0.2% TX-100 in PBS, following one night incubation in 4°C of primary antibodies, for which anti-CD63 antibody was used at a concentration of 1:200, meanwhile anti-vwf was 1:1000. Then secondary antibodies, concentration 1:500, conjugated to Alexa Fluor dyes.

### Granule release testing

The supernatant of activated platelet was collected and centrifuged at 4°C 1000×g for 15 min after stopping on the ice. Removing the supernatant to the eppendorf tube performed PF4 and 5-HT analysis according to the instructions of the Elisa Kit or reserved it in −80°C.

### Super-resolution microscopy of platelets

The image was acquired from structure illumination microscopy, one of the Superresolution microscopy applying.9 raw images taken by nine different sinusoidal illumination patterns were reconstructed to one final image waiting for analyzing[13]. Exposing time was 25ms. Sample was successively abrupted images under two different channel excitation light at 488nm (CD63) and 647 nm (vwf).

### Platelet morphology and ultrastructure distribution measurement

Platelet morphology was evaluated by average diameter calculated by Image J plugins in the rest stage. To assess the differences of average number in single platelet between patients and control groups, we calculated total number of CD63, vwf and platelet in each image using Image J, acquiring average number of CD63 and vwf of single platelet in each sample. The average size of CD63 and vwf was obtained from Image J. Mean area of CD63 and vwf per platelet was calculated by all quantity of CD63 or vwf over platelet number. %area of granules that means the percentage of CD63 or vwf in each image field was calculated by Image J.

### Data analysis

Data processing is anonymous. All the data was analysis by Graphpad Prism8.3.0. Unpaired, two-tailed student T-test was adopted to find statistically significant differences between patient and healthy control (healthy control that includes YH group and EH group) in platelet diameter, granule number, granule size, mean area of granule per platelet and %area of granule, if the data contented with normal distribution. Vwf average size was analysed by two-tailed Mann Whitney test because of the data doesn’t fit a normal distribution. P value <0.05 of T-test was regarded as significant differences.

## Results

### Significant changes in platelet morphology and ultrastructure were observed in AIS patients using GI-SIM

The CD63 and vwf structures were detected by immunofluorescence staining in dense granules and α granules of platelets, respectively. We observed morphology and ultrastructure of platelets in AIS and healthy control group with the help of SIM (Figure 1). Under SIM observation, the platelets of AIS patients were disc-shaped, with smooth and neat edges, and there was no obvious morphological change and difference compared with normal platelets (Fig 1). Dense granules and α granules are organelle structures in platelets with important function. It can be clearly seen that the two kinds of granules showed punctate distribution scattered in the platelet, with no aggregation between the granules by SIM (Fig 1). Subsequently, we observed the dense granules and α granules of platelets in AIS patients, and there was no obvious heterogeneity in morphology between AIS patients and healthy controls (Fig 1).

**Fig 1:**
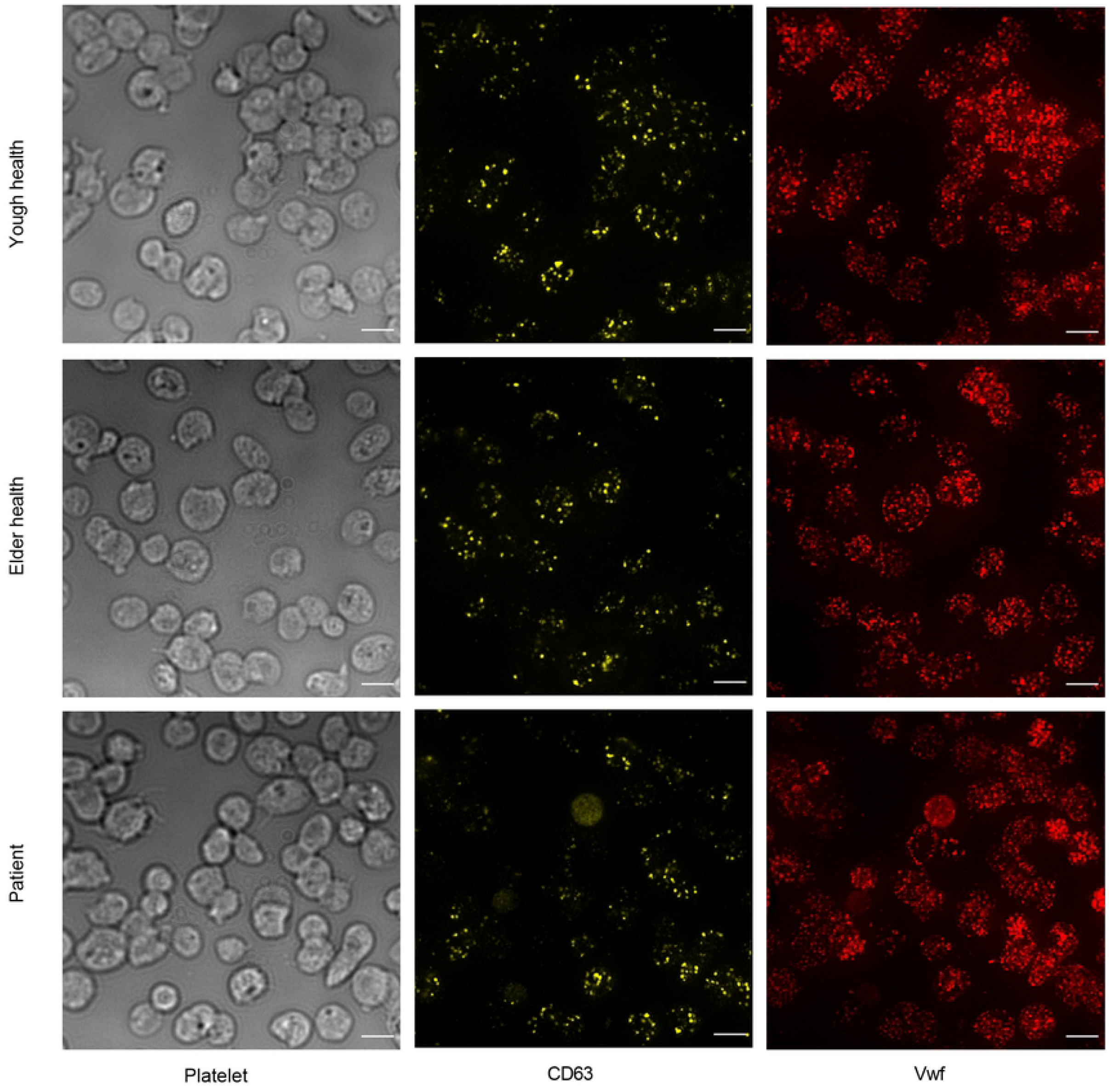
SIM shows morphology and ultrastructure of platelet in AIS patients and health group. A representative image of resting platelet is shown from young health (YH, n=6), elder health (EH, n=6) and AIS patient (n=7). The images show platelets are disc-shaped with smooth and neat edges in column 1; the dense granule and α granule labelled with anti-CD63 (yellow) and anti-vwf (red) in column 2 and column 3. The two-color images were acquired with laser light at 488 (CD63) and 642 (vwf). The data in this figure display oval shaped platelets and punctate distribution of dense granule and α granule scattered in the platelet. Scale bars: 3μm.

### Quantitative analysis of SIM images showed the diameter and average size of CD63 and vwf of single platelet reduced in AIS patients

Although no morphological differences were observed, platelet diameter was calculated from totally 1900 platelets in AIS patients and healthy subjects for comparison by SIM images (Fig 2a-b). In order to avoid the bias caused by the influence of age on platelet, we selected the elderly healthy group and the young healthy group as the healthy control group. Information on age, sex, NHISS score, MRI results, past history, medication history, platelet count, mean platelet distribution width, mean platelet volume, and ratio of large platelets between patients and controls are shown in Table 1 (Table 1). A total of 5372 platelets from 7 patients and 12 healthy controls (EH and YH) were selected. Finally, the average values of all samples were calculated. The mean platelet diameter of the patients was shorter than that of the elderly healthy group and the young healthy group (3.30±0.40 μm vs 3.60±0.29 μm, P=0.151; 3.30±0.40 μm vs 4.03±0.32 μm, P=0.004) (Figure 2b), and the results were significantly different. Otherwise, there were differences between elderly healthy group and the young healthy group (3.60±0.29 μm vs 4.03±0.32 μm, P=0.0356).

**Fig 2:**
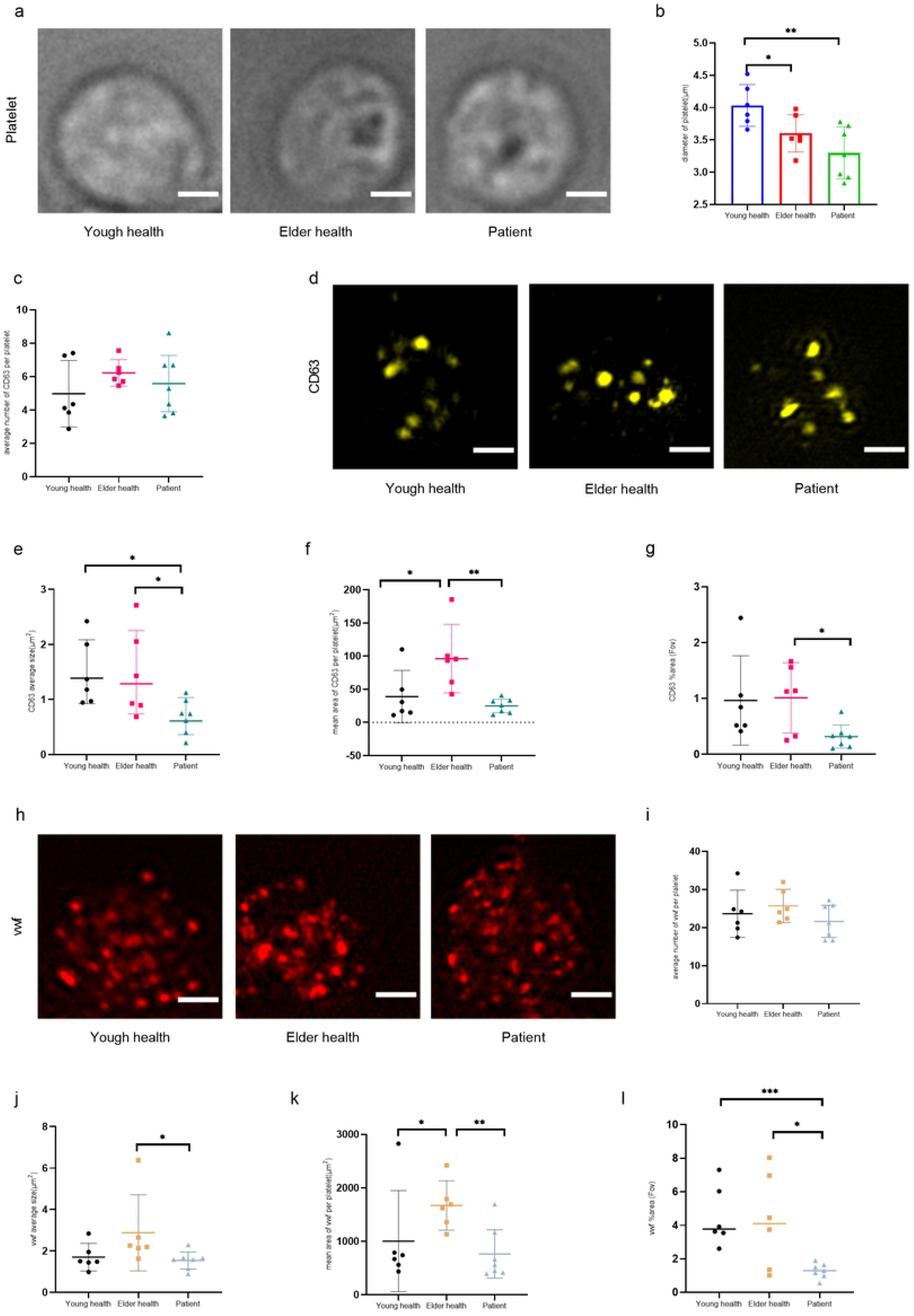
AIS patient has smaller size and area of CD63 and vwf. **a**, SIM imaging of platelets in young health (n=2521), elder health (n=2531) and AIS patient (n=2529). **b**, Platelet diameter decreased corelate with age. Comparison platelets diameter from YH (n=2521, 6YH), EH (n=2531, 6EH) and AIS patient (n=2529, 7 patients) showed platelets in elder health and AIS patient are smaller. Data are presented as mean with 95% CI. Unpaired two-tailed Student’s t-test. *P=0.0356, **P=0.0042. **c**, Counts of average number of CD63 in per platelet of YH (n=3143,6YH), EH (n=2921, 6EH) and AIS patient (n=3362, 7 patients) showed no difference. Unpaired two-tailed Student’s t-test. **d**, GI-SIM imaging of CD63 in young health (n=3143), elder health (n=2921) and AIS patient (n=3362). **e**, Quantification of average size of CD63 in YH, EH and AIS patient. CD63 average size in AIS patients was lower than YH (*P=0.038) and EH (*P=0.0111). **f**, Quantification of mean area of CD63 per platelet in YH, EH and AIS patient. Unpaired two-tailed Student’s t-test. *P=0.0467, **P=0.0032. **g**, Analysis of area% of CD63 field of view (Fov) in YH, EH and AIS patient. *P=0.0159. **h**, GI-SIM imaging of vwf in young health (n=3143), elder health (n=2921) and AIS patient (n=3362). **i**, Counts of average number of vwf in per platelet of YH (n=3143,6YH), EH (n=2921, 6EH) and AIS patient (n=3362, 7 patients) showed no difference. Unpaired two-tailed Student’s t-test. **j**, Quantification of average size of vwf in YH, EH and AIS patient. Mann-Whienty test used in EH vs AIS patient, unpaired two-tailed Student’s t-test in YH vs AIS patient and EH vs YH. vwf average size in AIS patients was lower than EH (*P=0.0111). **k**, Analysis of mean area of vwf per platelet in YH, EH and AIS patient. Unpaired two-tailed Student’s t-test in EH vs AIS patient, Kolmogorov-Smirnov test in YH vs EH. *P=0.026, **P=0.0051. **l**, Quantification of area% of CD63 field of view (Fov) in YH, EH and AIS patient. Unpaired two-tailed Student’s t-test. *P=0.0195, ***P = 0.0007. Scale bar:1μm.

**Table 1:** Characteristics of AIS patients and healthy control.

| Number | Sex | Age | NHIS score | MRI | Complication | Drug administration | Platelet number (125-350*10<sup>9</sup>) | PDW (15.5-18.1fL) | MPV(9.4-12.5fL) | P-LCR (17.5-24.3%) |
| --- | --- | --- | --- | --- | --- | --- | --- | --- | --- | --- |
| P1 | M | 58 | 0 | Lacunar infarction | hypertension<br>PU | / | 243 | 16.7 | 12.3 | 10.4 |
| P2 | F | 71 | 4 | Right basal ganglia infarction | hypertension | / | 234 | 14.1 | 11.2 | 35 |
| P3 | M | 59 | 3 | Coronal region infarction | hypertension<br>CKD | / | 232 | 12.9 | 11.1 | 33.8 |
| P4 | M | 54 | 6 | Right cerebellar infarction | hypertension<br>CKD | nifedipine | 158 | 16.1 | 12.2 | 43.4 |
| P5 | F | 85 | 3 | Left basal ganglia infarction | hypertension | / | 217 | 16.5 | 9.4 | 21.9 |
| P6 | M | 62 | 2 | Right parietal lobe infarction | / | / | 290 | 10.5 | 9.9 | 23.9 |
| P7 | M | 65 | 4 | Left parietal lobe infarction | / | / | 148 | 15.6 | 11.6 | 38.3 |
| EH1 | M | 74 | / | / | / | / | 223 | 12.4 | 10.5 | 29.3 |
| EH2 | F | 62 | / | / | PD | madopar | 203 | 13.2 | 11.7 | 37.8 |
| EH3 | M | 70 | / | / | APB | metoprolol | 288 | 10.9 | 9.9 | 24 |
| EH4 | M | 67 | / | / | / | / | 206 | 13 | 10.7 | 30 |
| EH5 | F | 65 | / | / | / | / | 163 | 17 | 13 | 48.3 |
| EH6 | M | 73 | / | / | / | / | 201 | 11 | 9.9 | 23.8 |
| YH1 | F | 24 | / | / | / | / | 253 | 11.1 | 8.9 | 18.5 |
| YH2 | M | 25 | / | / | / | / | 160 | 10 | 8.9 | 17.8 |
| YH3 | M | 22 | / | / | / | / | 224 | 10.7 | 9 | 18.9 |
| YH4 | M | 26 | / | / | / | / | 240 | 10.6 | 9.1 | 18.7 |
| YH5 | F | 21 | / | / | / | / | 209 | 8.5 | 7.9 | 19 |
| YH6 | M | 29 | / | / | / | / | 293 | 10.6 | 9.1 | 18.7 |

Patient(P), Elder health (EH), Young health (YH), National Institute of Health stroke scale(NIHSS), National Institute of Health stroke scale (MRI), Platelet distribution width(PDW), mean platelet volume(MPV), platelet large cell ratio (P-LCR).

Similarly, we quantitatively analyzed the image information of platelet dense granules and α granules obtained by SIM. Due to the heterogeneity of platelets, a total of 8171 platelets were selected and 8 parameters were measured: The number of CD63 and vwf, the average size of CD63 and vwf, the mean area of CD63 and vwf per platelet, as well as the CD63 and vwf area% in each field. In addition to the number of granules, the average size of platelet dense granules and α granules, the mean area of platelet within each platelet, and the area% in each field were significantly reduced in AIS patients (Fig 2). The CD63 average size of AIS patient was significantly lower than healthy control including EH and YH (0.69±0.32 μm^2^ vs 1.45±0.79 μm^2^, P=0.038; 0.69±0.32 μm^2^ vs 1.48±0.60 μm^2^, P=0.0111; Figure 2e). Meanwhile, there were no difference between EH and YH (1.45±0.79 μm^2^ vs 1.48±0.60 μm^2^, P=0.942, Fig 2e). The mean area of CD63 per platelet in AIS patients was significantly lower than that in EH group, but there was no difference between patient group and YH group (24.88±10.88 μm^2^ vs 96.20±49.16 μm^2^, P=0.0032; 24.88±10.88 μm^2^ vs 38.89±37.61 μm^2^, P=0.364, Fig 2f). In the meantime, this parameters in EH group was higher than YH group (96.20±49.16 μm^2^ vs 38.89±37.61 μm^2^, P=0.0467, Figure 2f). The CD63 area% in each field in AIS patient was smaller than healthy control including EH and YH (0.3±0.2 vs 1.0±0.6, P=0.0159; 0.3±0.2 vs 0.9±0.7, P=0.0541; Fig 2g). When assessing vwf, the other ultrastructure of platelets, there were similar results like CD63. Firstly, the average size of vwf in AIS patient was also lower than healthy control including EH and YH (1.54±0.44 μm^2^ vs 2.87±1.75 μm^2^, P=0.035; 1.54±0.44 μm^2^ vs 1.70±0.64 μm^2^, P=0.6084; Fig 2j). Meanwhile, there were no difference between EH and YH (2.87±1.75 μm^2^ vs 1.48±1.70±0.64 μm^2^, P=0.0649, Fig 2j). Then, the mean area of vwf per platelet in AIS patients was lower than that in EH group and YH group, and there was significantly difference between patient group and EH group (765.03±488.57 μm^2^ vs 1670.91±439.53 μm^2^, P=0.0051; 765.03±488.57 μm^2^ vs 1004.78±902.08 μm^2^, P=0.4452, Fig 2k). In the meantime, this parameters in EH group was higher than YH group but with no difference (1670.91±439.53 μm^2^ vs 1004.78±902.08 μm^2^, P=0.0649, Fig 2k). Finally, the vwf area% in each field in AIS patient was smaller than healthy control including EH and YH (1.29±0.44 vs 4.26±2.86, P=0.0195; 1.29±0.44 vs 4.51±1.78, P=0.0007; Fig 2l). The other two parameters of CD63 number and vwf number had no statistical difference (Fig 2c, i).

### α granule in platelet of AIS patients changed to a pattern of parenchymatous masses upon stimulation of TH observed by GI-SIM

Platelets were stimulated with 0.5U thrombin (TH) in healthy subjects and AIS patients, and the morphological changes of platelets, platelet dense granules and α granules were observed by SIM at different time periods of 0min, 5min and 1h (Fig 3a). 5 minutes after activation, platelets gradually spread out from the disk and extending pseudopodia. α granules began to aggregate and adhere to each other in both healthy people and AIS patients (Fig 3a, S1). Multiple platelets gathered in the same area to form blood clots, and the shape of a single platelet could not be recognized after 1h activation (Fig 3a). At this time point, α granules showed morphological differences between AIS patients and healthy people. In AIS patients, α-granules accumulate together to form large fluorophores with few diffuse particles around them (Fig 3c). The formation of α granules with less gap was closer to the structure of parenchymatous masses. However, in healthy people, no matter EH or YH, there are more gaps between α granules to form areolar structure masses, and more granular parts are scattered around the masses (Fig 3a, c). There was no significant difference in the morphology of dense granules at 0min, 5min and 1h after thrombin activation (Fig 3b).

**Figure 3:**
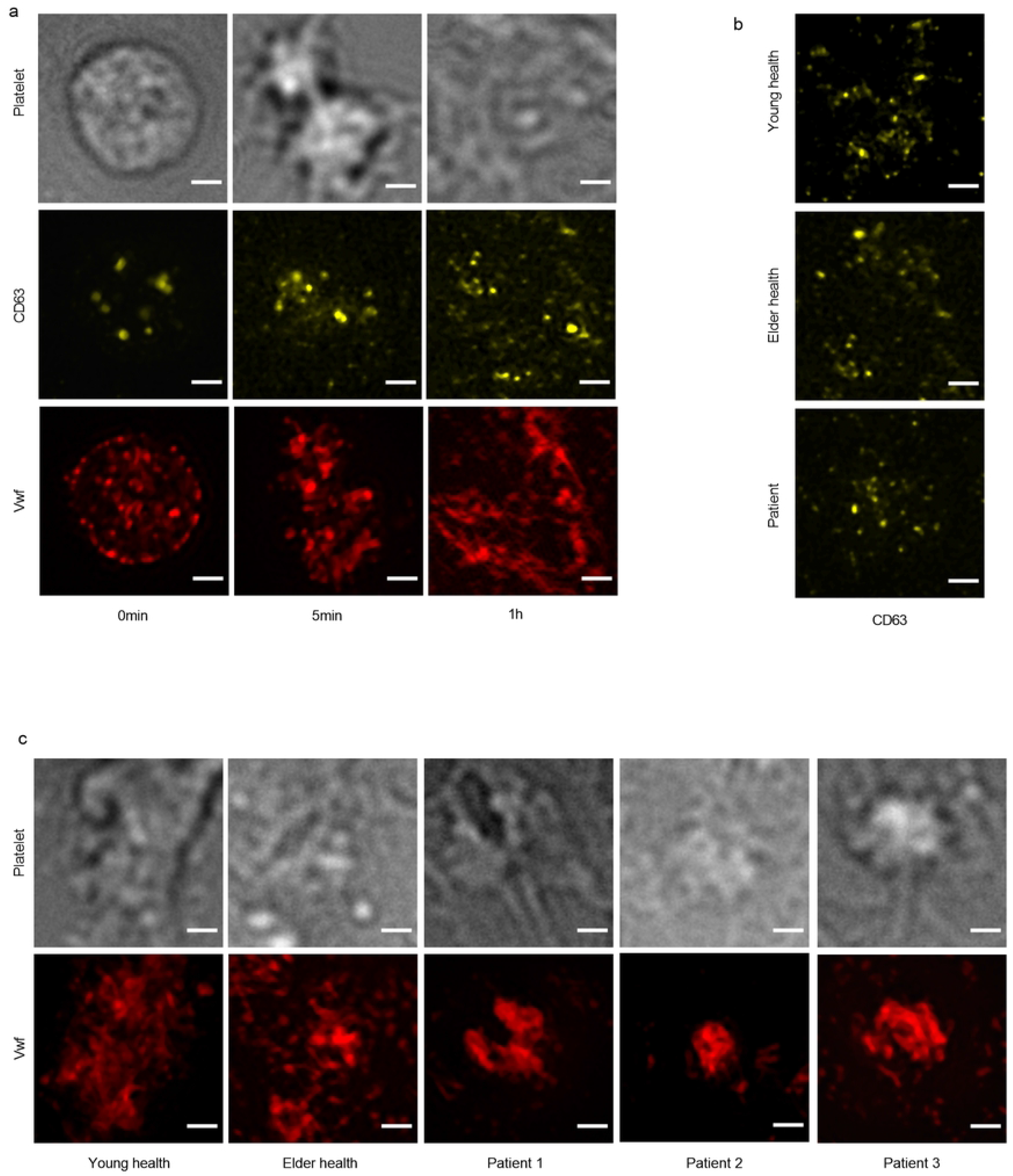
α granule displayed a pattern of parenchymatous fluorophores masses under stimulation in AIS patients. a, SIM images of platelet activated by 0.5U TH after 0min, 5min and 1h from one health control (n=12). CD63(yellow) showed no significantly difference in morphology at different time point. Vwf (red) aggregation increased over time and presented areolar structure masses in 1h. b, CD63 showed no difference between YH (n=6), EH (n=6) and AIS patient (n=7) after 1h stimulation. c, vwf altered to parenchymatous fluorophores masses in AIS patients(n=7) while health people (n=12) presented areolar structure masses after 1h stimulation. Scale bar:1μm.

### 2MeSamp inhibited the formation of parenchymatous masses morphology of platelet α granules in AIS patients

2MeSAMP and Aspirin are two kinds of antiplatelet drugs that inhibit ADP receptor and COX-1 receptor, respectively. Platelets of AIS patients were treated with 2MeSAMP and Aspirin at concentrations of 10μM and 53μM in vitro, respectively, and then activated with 0.5U TH for 1h. More single platelets could be observed after treated by 2MeSAMP which belongs to ADP antagonist in the wide field, compared with direct or aspirin treatment (S2). In the meantime, the α granules of platelets in AIS patients changed from parenchymatous masses morphology to areolar structure masses after treatment with 2MeSAMP (Fig 4a). On the other hand, this alteration hasn’t been observed after aspirin treated (Fig 4a). This phenomenon showed that 2MeSAMP inhibit the formation of parenchymatous masses morphology of platelet α granules in AIS patients. And it indicated that parenchymatous masses morphology of α granules in platelets may represent a higher level of activation. No morphological changes were observed in dense granules (Fi4b, S3).

**Fig 4:**
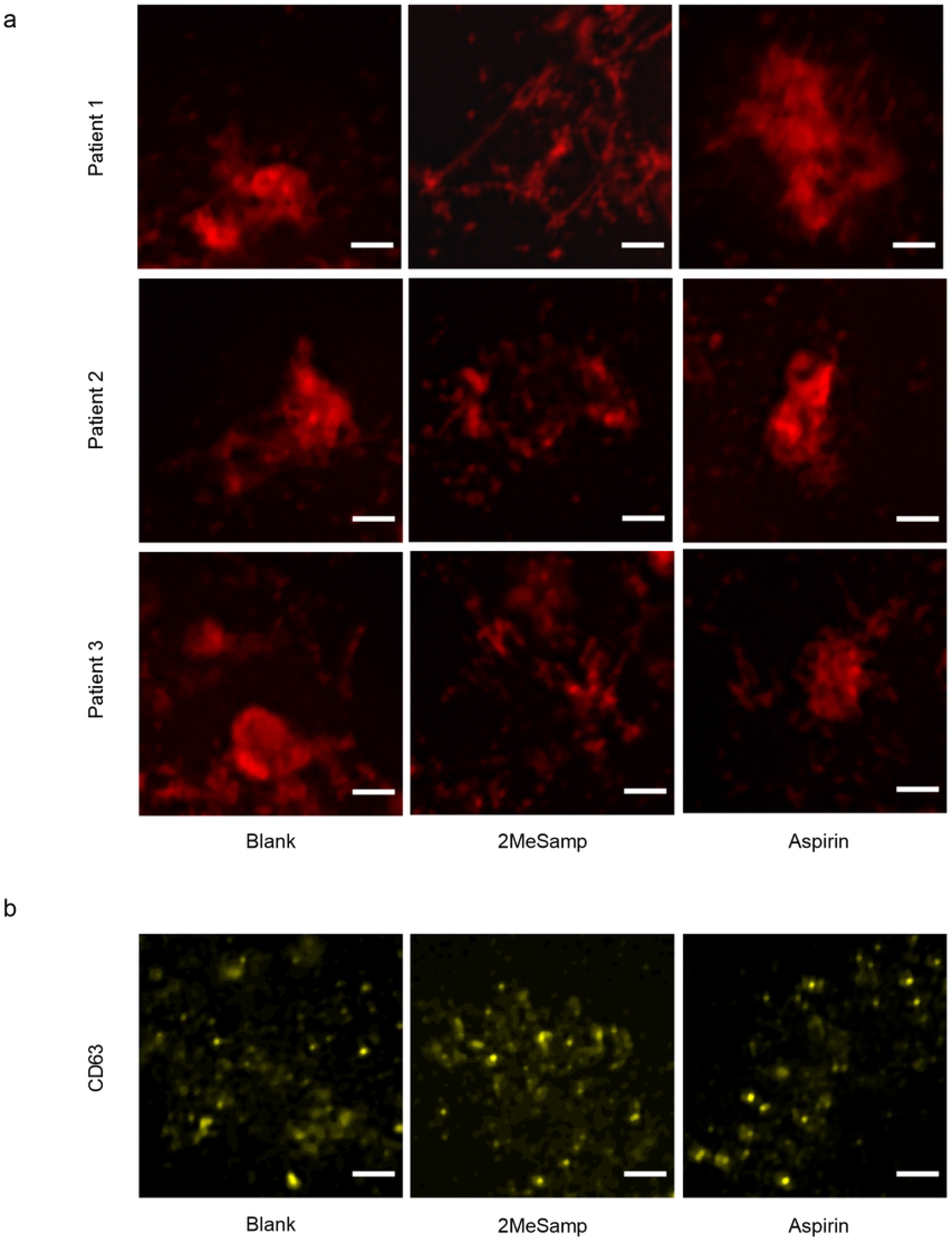
2MeSamp depolymerized α-granule fluorophores. a, SIM image of platelet vwf (red) from 3 of 7 AIS patients. Under 0.5U TH stimulation, no drugs administrated in column 1, treated with 2MeSamp in column 2, treated with aspirin in column 3. The results showed 2MeSamp depolymerized parenchymatous fluorophores masses of α granule. And aspirin didn’t influence the fluorophores formation of α granule. b, platelet CD63 (yellow) morphology unchanged under different treatment. Scale bar: 1 μm.

### Release level of PF4 was higher in AIS patient upon TH stimulation and treatment with 2MeSamp inhibited PF4 releasing

PF4 is a protein released by α granules. We measured the release of PF4 to evaluate the the function of α granules between AIS patients and healthy controls. The levels of PF4 in EH were 233.95±4.57ng/ml, 493.77±16.27ng/ml and 1992.25±57.77ng/ml at rest, 5min after activation and 1h after activation, respectively (Fig 5a). The levels of PF4 in AIS patients were 147.15±2.51ng/ml, 468.82±27.02g/ml and 1272.48±80.91ng/ml at rest, 5min after activation and 1h after activation, respectively (Fig 5b). The PF4 releasing of AIS patient was significantly lower than healthy group in rest stage (147.15±2.51ng/ml vs 233.95±4.57ng/ml, P<0.0001; Fig 5c). Due to the heterogeneity of platelets, it is not reasonable to directly compare the release of PF4 from α granules in platelets with AIS patients and healthy controls. Therefore, the platelet release from AIS patients and healthy controls were normalized and then compared. The fold change of PF4 release after 5min and 1h activation in patients and healthy subjects were (3.19±0.18 vs 2.11±0.07, P=0.0007; Figure 5d) and (8.65±0.55 vs 8.52±0.25, P=0.724; Fig 5d). The release of PF4 from platelets in AIS patients after treatment with 2MeSamp and Aspirin in vitro and the ratio of PF4 release with direct activation can reflect the effect of the inhibitor. Regardless of 5min or 1h activation, 2MeSamp had a greater inhibitory effect on PF4 release than Aspirin (0.67±0.09 vs 1.67±0.19, P=0.0012; 0.57 ± 0.08 vs 0.02±0.87, P = 0.003; Fig 5f).

**Fig 5:**
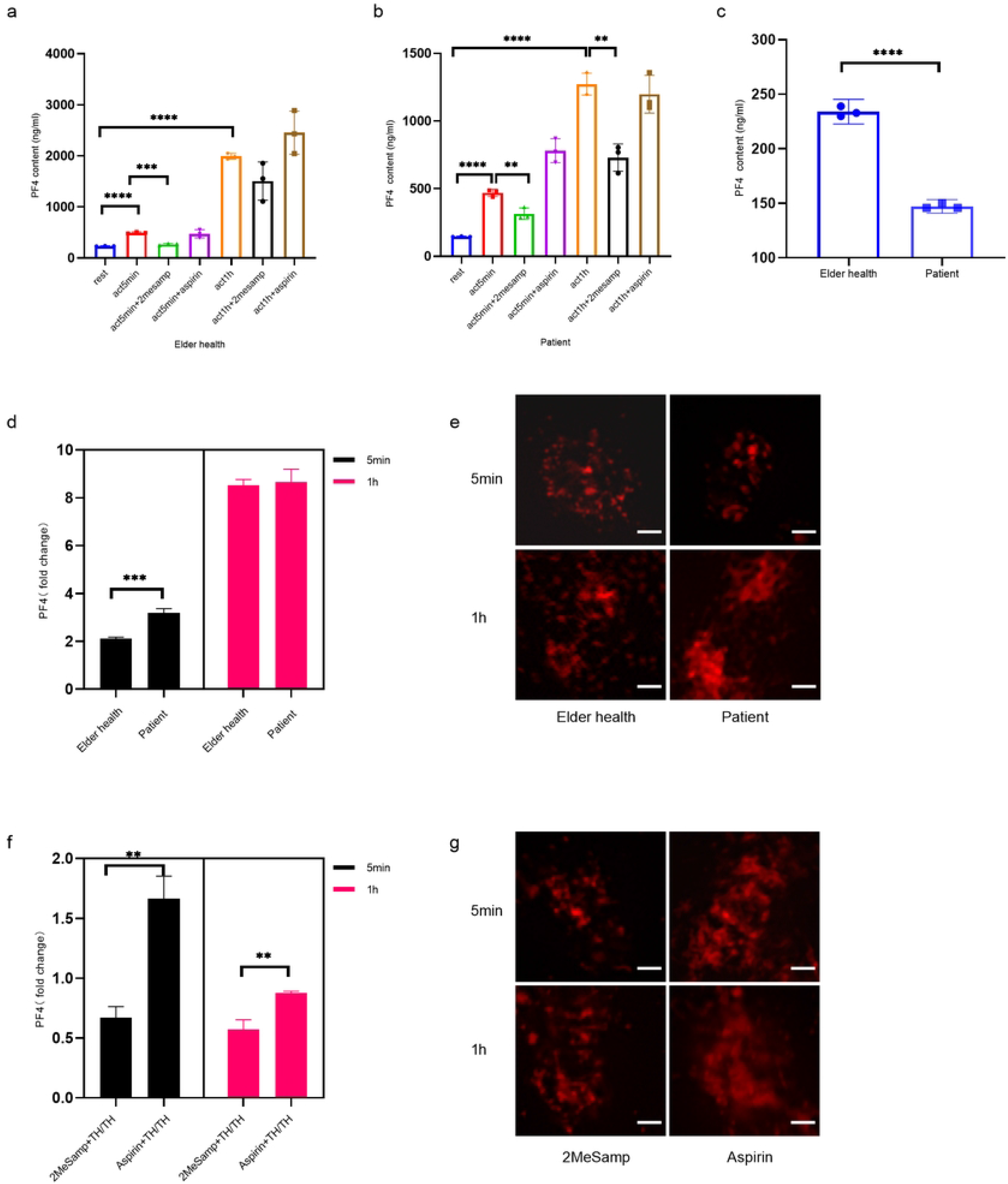
PF4 content was lower in AIS patient but release more after being stimulated while 2MeSamp inhibited PF4 releasing. **a,** PF4 content at 7 different treatments (rest; activate for 5min; dealt with 2MeSamp and activate for 5min; dealt with Aspirin and activate for 5min; activate for 1h; dealt with 2MeSamp and activate for 1h; dealt with Aspirin and activate for 1h) in EH (n=3). Data are presented as mean with 95% CI. Unpaired two-tailed Student’s t-test. PF4 content increased after activating 5min and 1h compared with rest stage (****P <0.0001; ****P <0.0001). Releasing of PF4 decreased after dealing with 2MeSamp and activating for 5min and 1h (***P =0.0001). **b**, PF4 content at 7 different treatments in AIS patients (n=3). Unpaired two-tailed Student’s t-test. PF4 content increased after activating 5min and 1h compared with rest stage (****P <0.0001; ****P <0.0001). Releasing of PF4 decreased after dealing with 2MeSamp and activating for 5min and 1h (**P =0.0062; **P =0.0019). **c**, PF4 content was lower in AIS patients(n=3) at rest stage (****P <0.0001). Unpaired two-tailed Student’s t-test. **d**, Under stimulation with 0.5U TH for 5min, the rising multiple was higher in AIS patients (n=3) compared with EH (n=3) (***P=0.0007). Unpaired two-tailed Student’s t-test. **e**, GI-SIM image of vwf in activated platelets at 5min and 1h from one EH (n=6) and one AIS patient (n=7). **f**, Treatment with 2MeSamp inhibited the release of PF4 after 5min and 1h stimulation compared with aspirin (**P =0.0012; **P =0.003). **g**, Imaging of vwf activated 5min and 1h from one patient (n=7) after treatment with 2MeSamp and Aspirin respectively. Scale bar: 1μm.

## Discussion

Electron microscopy can see cells at the single-nanometre level, but sample preparation is time-consuming and requires experienced researchers to identify specific organelles. SIM, a new imaging technique, employs immunofluorescence to label the subcellular structure of cells, which has the advantages of high resolution, simple sample production, short time consuming, less damage to samples and easier imaging. The advantages of SIM enable us to better examine the morphology and ultrastructure of platelets in AIS patients. It is easier to obtain more accurate picture information, including platelet morphology, the structure of platelet dense granules and α-granules. At the same time, we obtain a large number of picture information to process and conduct parameter analysis.

Platelets from healthy controls were in a resting state because they had a smooth surface and were free of pseudopodia and aggregation, which are considered indicative of platelet activation. There was no obvious heterogeneity in platelet morphology between the patient group and the healthy control group. However, the mean platelet diameter of AIS patients and EH group was smaller than that of YH group, which means that the mean platelet diameter became smaller with age. Previous studies of platelets from AIS or TIA patients by electron microscopy have shown that platelets from AIS or TIA patients have more pseudopodia and spread than those from healthy subjects[22]. We did not find this difference when we used SIM to study platelets in AIS patients. The reason may be that compared with electron microscopy, the sample preparation of SIM is easier and thus has less damage to platelets, which keeps platelets in the unstimulated resting state or the stimulation is too slight to cause platelet deformation. The difference of platelet morphology between SIM and electron microscopy also indicates that the platelet in AIS patients is not in the activated state at rest, but more likely to be at a hypersensitive state. This means platelets in AIS patients were more likely to show a large number of pseudopodia and aggregation activation after receiving the same stimulation that exceed the threshold stimulation.

A large number of subcellular structures exist in platelets. α granules and dense granules are absolutely necessary structures for platelet function after activation. We mainly analyzed the number, size and area of the granules in platelets. In addition to the number, the average size, area in single platelets and the area% of dense granules and α granules in AIS patients’ platelets were lower than those in healthy controls. As a consequence, previous imaging methods were relatively imprecise for the reason that they can only count the number of granules, lacking the evaluation of the size and area of dense particles and α particles. The high resolution of SIM makes it possible to process the image information to obtain the corresponding parameters of the size and area of platelet dense granules and α granules. Therefore, with the application of SIM, we can objectively analyze the structural changes of the two kinds of granules in AIS, especially when the granule number have no difference. Platelets’ differences founded by SIM in AIS patients may inform we could consider testing ultrastructure of platelets to find out people with abnormalities earlier who may have a high-risk in acquiring an AIS. We could reduce the risk of AIS in these populations through early and appropriate intervention.

In AIS patients, the size and area of platelet dense granules and α granules were decreased compared with healthy controls (Fig 2). The mechanism of this change may be due to the highly sensitive state of circulating platelets in AIS patients, and the body initiate negative feedback regulation to reduce the abnormal activation of platelets. Therefore, in AIS patients, the number of platelet dense granules and α granules has not changed significantly, but the structure has changed in response to the hypersensitive state of platelets. As mentioned before, platelets in AIS patients are more likely to be activated in response to supramental stimulation which means they are in a highly sensitive state. This was confirmed in functional assays of platelet alpha granules, in which fold change of PF4 release were significantly higher in AIS patients than in healthy controls at 5min of activation under equal conditions (Fig 5d).

When 0.5U thrombin was used to activate platelets in AIS patients and healthy controls, it was found that platelet α granules in AIS patients changed from dispersed round granules to a pattern of parenchymatous masses (Fig 3c). However, the α-granules in the healthy control group showed areolar structure like masses (Fig 3a). The change of α granule from scattered granular to massive may be the aggregation of multiple granules, which can be regarded as a sign of platelet activation. After treatment with the antiplatelet drug 2MeSAMP, platelets in AIS patients were activated with the same concentration of thrombin, and α granules changed from parenchymatous masses to areolar structure masses (Fig 4a). Therefore, we believe that the parenchymatous masses of α granules is a manifestation of greater activation compared with the areolar structure masses. Similarly, parenchyma mass also indicates that platelets in AIS patients are in a hypersensitive state. When subjected to the same degree of stimulation, the platelets of AIS patients are more likely to activate and α granules are more closely bound. α granules contain hundreds of proteins, including vwf, fibrinogen, and numerous coagulation factors. Among them, vwf and fibrinogen are two adhesion proteins, which play an important role in platelet aggregation and adhesion to the damaged site [2]. Therefore, the aggregation of platelet α granules in AIS patients was more tightly clumped, indicating that platelets in AIS patients may play a more powerful role in aggregation and adhesion after activation than healthy people. This is also consistent with the conclusion that platelets in AIS patients are hyperactivated in previous studies [23, 24].

Platelets from AIS patients were treated with 2MeSAMP, an ADP receptor antagonist with in vitro activity, and then activated with 0.5U thrombin. Platelets α granules dispersed from parenchyma to areolar structure masses (Fig 4a). In contrast, treatment with aspirin that is a COX-1 inhibitor showed no significant change in platelet α granules. We considered that this may represent a stronger inhibitory effect of 2MeSAMP on platelets, in the absence of metabolic effects in vivo. The release of PF4 from α granule was in deed significantly lower after treatment with 2MeSAMP than aspirin treatment (Fig 5f). 2MeSAMP has the same mechanism of action as clopidogrel, an antiplatelet drug commonly used in clinic, acting through inhibiting ADP binding to P2Y12 receptor. Some clinical trials have also demonstrated that clopidogrel is more effective than aspirin in antiplatelet effects in AIS patients [25, 26]. Combined with our experimental results, the reason for the stronger inhibitory effect of clopidogrel on platelets may be that it not only inhibits P2Y12 receptor, but also affects the release of α granule.

Our study found that platelet in AIS patient was in a hyperactive stage. There were significantly decrease in average size, area% of dense granule and α granule as well as mean area of dense and α granule per platelet. These alterations indicated that ultrastructure in platelet has changed during AIS processing. Moreover, α granule in platelets from AIS patients congregated to center and changed to a pattern of parenchymatous fluorescent masses after being stimulated, which indicated a stronger activated state. The reduction of granule size, mean area of granules per platelet and area% in resting platelets of AIS patients may represent a negative feedback regulation mechanism. This can be proofed by lower PF4 release in AIS patients which indicated that the function of α granules were inhibited in rest state (Fig 4c). However, the platelets of AIS patients present a hyperactivity after stimulating (Fig 4d). Upon stimulation of TH, different morphology of α granule in platelet of AIS patients and healthy control might correlation with the hyperresponsiveness of platelets in AIS patient as α granule masses depolymerized after dealing with antiplatelet drug 2MeSAMP. Furthermore, more tightly gathering of α granule probably relative to increasing fibrin fiber thickness which has been demonstrated that the platelets of TIA patient formed thicker fibrin fiber after being stimulated[22].

Otherwise, 2MeSAMP has stronger depolymerization to α granule masses in activated platelets than aspirin (Fig 4a). Combined with release test of granule, 2MeSAMP in deed inhibited granule releasing function in some extent contrasting with aspirin(Fig 5f). We can conclude that 2MeSAMP not only suppresses the combination of P2Y12 receptor, but also change the ultrastructure and function of α granule in activated platelets. Several studies have demonstrated that P2Y12 receptor has an influence on granule release, fibrinogen receptor activation and TXA2 production besides function on ADP receptor[27, 28]. As a consequence, antagonizing P2Y12 receptor means exerting antiplatelet effects in a variety of ways which may show a better efficacy of antiplatelet.

In view of 2MeSAMP belongs to P2Y12 receptor antagonist, this may indicate that P2Y12 receptor antagonist is more effective than aspirin which is a common COX-1 inhibitor in vitro combining our results. A number of clinical studies have given opinion that P2Y12 receptor antagonist was more effective in protecting patients with AIS or coronary heart disease than COX-1 inhibitor. A retrospective cohort study by Hsin-Yi et al. showed that clopidogrel could significantly reduce recurrent stroke compared with aspirin[29]. Maurizio et al. draw a similar conclusion that clopidogrel is better in clinical benefit to AIS patient than aspirin[30]. However, there are some studies have a contrary opinion that clopidogrel showed a higher risk of recurrent stroke in comparison with aspirin[31]. This variant may result from antiplatelet drug resistance, racial difference and individual differences of patients. In virtue of SIM, we observed more obvious ultrastructure change in platelets after treatment of 2MeSAMP than aspirin, which indicate antagonize P2Y12 receptor may have a greater influences in inhibiting platelets activation compared to blocking COX-1 and supply a new perspective in assessing the effectiveness of antiplatelet drug the same time.

## Conclusion

Although platelets play an important role in AIS disease, there is still thirsty of knowledge on platelet ultrastructure and its change in patients with AIS. Meanwhile, whether choice P2Y12 receptor antagonist or COX-1 inhibitor in the later stages of AIS is still controversial even though dual antiplatelet therapy has been recommended clearly in AIS acute stage. On the strengthen of SIM, we found the ultrastructure of resting platelets in AIS patients is heterogeneous in granule size, mean area of granule in per platelet and granule area% compared with healthy people. And the granule release is lower in AIS patient in rest stage. Under activated circumstance, α granule in platelets from AIS patients congregated to a pattern of parenchymatous fluorescent masses. Otherwise, 2MeSAMP has stronger depolymerization to α granule masses in activated platelets than aspirin. These results indicated that platelets activity in AIS patients were negative in rest stage and alter to hyperresponsiveness after being stimulated. In the future, we may use SIM to detect the platelets in high-risk population of AIS. And P2Y12 receptor antagonist could be considered as a more effective antiplatelet drug for long-term monotherapy. Our study makes up for the lack of understanding of platelet ultrastructure in AIS patients. Meanwhile, platelets testing based on SIM may provide a new method of finding high-stake AIS people, diagnosing and evaluating efficacy of antiplatelet drug in AIS patients.

## Abbreviation

ADP: adenosine diphosphate
AIS: acute ischemic stroke
β-TG: β-thromboglobulin
CVD: cerebrovascular disease
COX-1: cyclooxygenase-1
EH: elder healthy people (>60 years old)
PF4: platelet factor 4
SIM: super-resolution microscope
TH: thrombin
TIA: transient ischemic attacks
vwf: von willebrand factor
YH: young healthy people (18~60 years old)
2MeSAMP: 2-methyl-thioadenosine-monophosphate, in vivo form of p2y12 receptor antagonist.

## Acknowledgements

We thank all the volunteers for their participation in the study. The authors thank Huan Deng and Qian Jiang for assisting in laboratory equipment. And we thank all other staff at neurology department in Wuhan Puai Hospital for help collecting samples, as well as blood transfusion laboratory of Wuhan Blood Center for supplying equipments.

**Correspondence** and requests for materials should be addressed to Bingxin Yang.

**S1: Platelet ultrastructure after 5min stimulation.**

**S2: 2MeSamp inhibit platelet aggregate.**

**S3: Antiplatelet drug didn’t influence morphology of CD63 in activated platelets.**

